# Paired high-throughput, in-situ imaging and high-throughput sequencing illuminate acantharian abundance and vertical distribution

**DOI:** 10.1101/2020.02.27.967349

**Authors:** Margaret Mars Brisbin, Otis Davey Brunner, Mary Matilda Grossmann, Satoshi Mitarai

## Abstract

Acantharians are important contributors to surface primary production and to carbon flux to the deep sea, but are often underestimated because their delicate structures are destroyed by plankton nets or dissolved by preservatives. As a result, relatively little is known about acantharian biology, especially regarding their life cycles. Here, we take a paired approach, bringing together high-throughput, in-situ imaging and high-throughput sequencing, to investigate acantharian abundance, vertical distribution, and life-history in the western North Pacific. Observed concentrations of acantharian cells correlated well with sequence abundances from acantharians with known, recognizable morphologies, but not to sequences from those without known morphology (basal environmental clades). These results suggest basal clades may lack characteristic star-shaped skeletons or are much smaller than known acantharians. The decreased size-range of acantharians imaged at depth supports current hypotheses regarding asymbiotic acantharian life cycles: cysts or vegetative cells release reproductive swarmer cells at depth and juvenile cells grow as they ascend towards the surface. Moreover, sequencing data present the possibility that photosymbiotic acantharians also reproduce at depth, like their asymbiotic, encysting relatives, which is counter to previous hypotheses. Finally, in-situ imaging captured a new acantharian behavior that may be a previously undescribed predation strategy.

## Introduction

Acantharians are important contributors to primary production in surface waters and to carbon flux to the deep sea (Michaels, 1991; Michaels et al., 1995; Decelle et al., 2013; Belcher et al., 2018). Clade E and F acantharians—the majority of described acantharian species—host algal endosymbionts from the haptophyte genus *Phaeocystis* (Decelle et al., 2012a; Mars Brisbin et al., 2018), which exhibit elevated photosynthetic efficiency when living symbiotically (Decelle et al., 2019). The acantharian skeleton is composed of strontium sulfate, the densest known organic biomineral, causing acantharians to sink quickly after death (Decelle et al., 2013). In addition, their amoeboid cell structure with sticky cellular extensions (pseudopodia) predisposes sinking acantharians to form aggregates, further enhancing sinking rate (Gutierrez-Rodriguez et al., 2019). The biogeochemical significance of acantharians has been historically underestimated, however, because traditional sampling methods often miss acantharians; plankton nets destroy delicate acantharian cell structures (Michaels, 1988) and common preservatives dissolve acantharian skeletons (Bernstein et al., 1992). DNA sequencing surveys have revealed that acantharians account for large numbers of sequences from the water column and sediment traps in diverse ecosystems, including tropical and subtropical regions (Fontanez et al., 2015; Hu et al., 2018), polar regions (Martin et al., 2010; Decelle et al., 2013), and productive temperate coastal regions (Countway et al., 2010; Gutierrez-Rodriguez et al., 2019). However, the relationship between DNA sequence abundance and acantharian biomass or flux is not clear and is complicated by acantharians being multinucleated and having multiple life stages, including encystment and reproductive swarmer production.

DNA metabarcoding—sequencing a region of the small-subunit ribosomal RNA gene for an entire community—has been extensively applied to estimating microbial community structure (e.g. (de Vargas et al., 2015; Pernice et al., 2016). While this method has undoubtedly revolutionized our understanding of microbial diversity in different ecosystems, it has several significant limitations. First, metabarcoding and other meta ‘omics produce compositional data, meaning that the abundance of any single group is inherently influenced by the abundance of other groups (Gloor et al., 2017). This issue is further complicated by the varying nucleus and gene-copy numbers among protist groups—some organisms, like dinoflagellates, have many copies of ribosomal RNA genes and may be more represented in sequencing datasets as a result (Gong et al., 2013; Gong and Marchetti, 2019). Gene-copy and nucleus number is especially problematic because it precludes the possibility of extrapolating absolute abundances from sequencing reads and total cell counts—if the cell abundance to sequence abundance ratio was consistent, absolute abundance could be determined by multiplying the relative read abundance with the total cell count in a sample (Gong and Marchetti, 2019). The second major limitation of DNA metabarcoding is that DNA can persist after a cell dies and, therefore, does not reflect metabolic state. As a result, it is unknowable whether DNA sequencing reads derive from actively metabolizing cells, dormant cells or cysts, reproductive cells, or dead cells and detritus (Torti et al., 2015). This is particularly relevant in evaluating acantharian abundances and relative contributions to biogeochemical cycles since acantharian vegetative cells, reproductive cells, and cysts will differentially contribute to photosynthesis, grazing/predation, and carbon flux (Decelle et al., 2013).

While not yet as widely adopted as molecular methods, high-throughput, in-situ imaging systems are being used to quantitatively assess abundances of marine microbes and other components of the plankton (Dennett et al., 2002; Grossmann et al., 2015; Biard et al., 2016). Such imaging systems can drastically improve the spatial resolution of sampling and process much larger volumes of water than are included in DNA surveys. Furthermore, imaging cells where they naturally occur and in their native orientation can reveal previously undescribed behaviors and associations (Möller et al., 2012; Greer et al., 2013; Peacock et al., 2014). Analyzing data from high-throughput imaging, however, is still challenging; processing images and creating training sets for use with machine learning algorithms requires expertise in plankton taxonomy and is time-intensive (Orenstein et al., 2015). The taxonomic resolution attainable with a particular imaging system depends on image size and quality, but will almost always be less than is possible with molecular methods. Furthermore, taxonomic resolution will vary for different taxonomic groups and will be higher for those with more defined morphological features and lower for organisms, like flagellates, that lack identifying features (Sieracki et al., 2010). Finally, a single imaging system cannot image the entire size-range of marine plankton, necessitating multiple systems to holistically characterize plankton communities (Lombard et al., 2019). Vegetative acantharian cells, with their characteristic star-shaped skeletons, are particularly amenable to imaging surveys (Biard et al., 2016), but distinguishing acantharian reproductive cells or cysts with high-throughput, in-situ imaging is probably not possible.

DNA metabarcoding and high-throughput, in-situ imaging both have benefits and drawbacks as methods for assessing plankton abundance and community structure. By applying these methods together, we aim to better characterize acantharian abundance, water-column distribution, and life-history. In this study, we deployed the BellaMare In-Situ Ichthyoplankton Imaging System (ISIIS) small-imager/area-scanner (Cowen and Guigand, 2008; www.planktonimaging.com/smaller-imagers) at four sites along the Ryukyu Archipelago in the western North Pacific. We additionally collected replicate water samples for DNA sequencing from the surface, subsurface chlorophyll maximum, middle water column, and about 10 m above the seafloor from each site where imaging was performed and 10 additional sites along the Ryukyu Archipelago. Water samples were sequentially size-fractionated in an effort to separate acantharian vegetative cells and cysts from reproductive swarmers. We compare the relative abundance of acantharian reads in the larger size fraction with cell counts from imaging profiles to assess the relationship between acantharian relative read abundance and cell abundance. Metabarcoding results are further analyzed to evaluate the taxonomic distribution of acantharians by depth in the western North Pacific and we consider results from size fractionation in the context of hypothesized acantharian life-cycles.

## Methods

### Sampling Locations

Water samples for DNA sequencing were collected from 14 sites spanning the length of the Ryukyu Archipelago during the Japan Agency for Marine-Earth Science and Technology (JAMSTEC) MR17-03C cruise from May 29 to June 13, 2017 (Fig. 1). The JAMSTEC DEEP TOW 6KCTD system, a towable frame outfitted with several imaging systems and a Conductivity-Temperature-Depth (CTD) sensor, was additionally deployed at four of the sampling sites (3, 10, 15, and 17), to take vertical profiles of plankton images from the sea surface to a maximum of 1,000 m (Fig. S1A).

**Fig 1.**
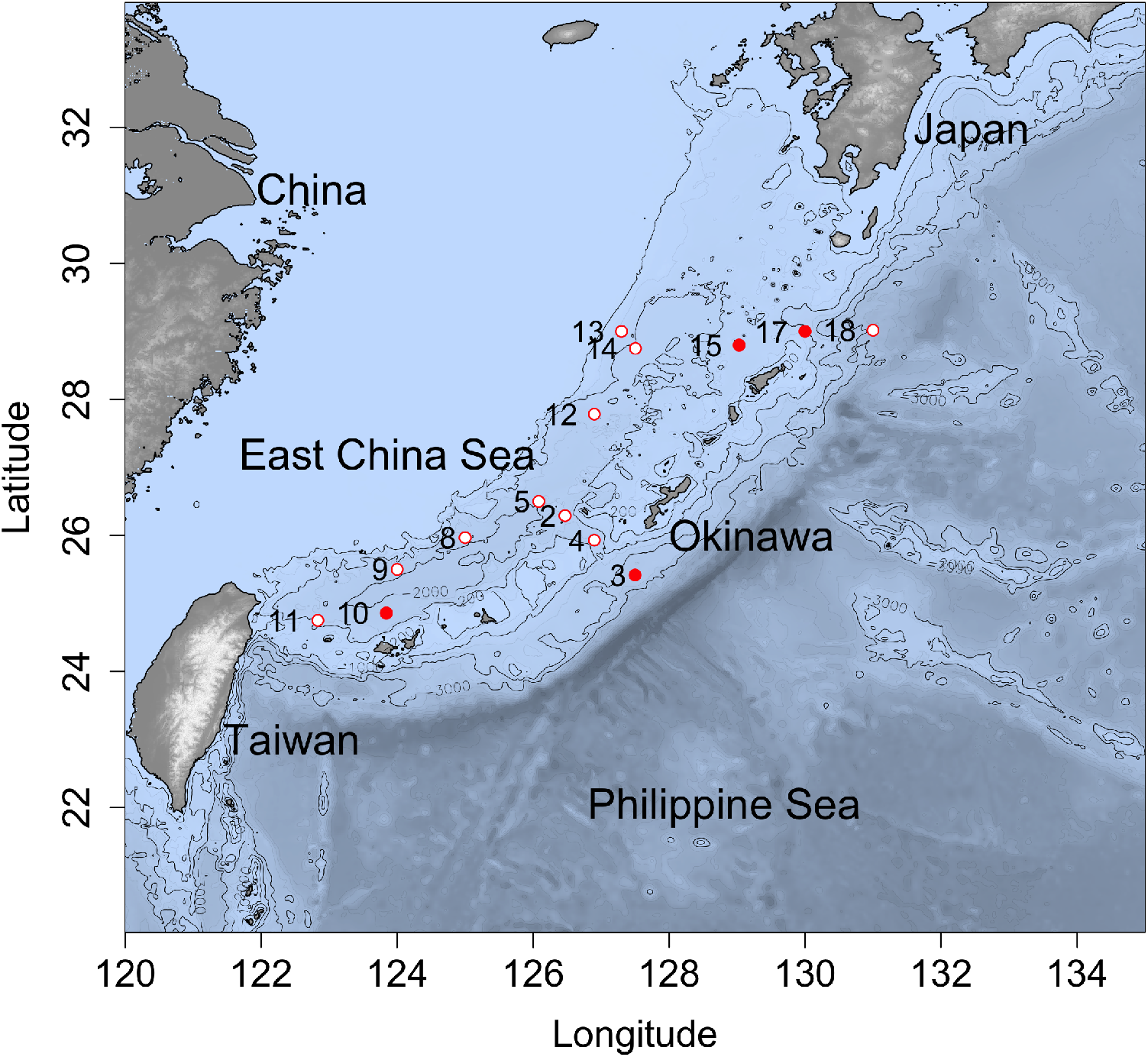
Map of sampling station locations. Stations where water samples for DNA analysis were collected and high-throughput imaging was performed are marked with closed red circles. Stations where only water samples for DNA analysis were collected are marked with open red circles.

### Image Acquisition & Processing

An ISIIS small imager/area-scanner (BellaMare, San Diego, CA) was attached to the DEEP TOW to collect vertical profiles of plankton images (Fig. S1). The camera system was set to image organisms > 250 μm, which aligns well with the size of vegetative acantharian cells. The ISIIS camera was programmed to take 1 photo per second coinciding with an LED flash. Each photo imaged 0.39 L (st. 3 and 10) or 0.35 L (st. 15 and 17) parcels of water in 2448 × 2050 pixel resolution, with each pixel being 22.5 × 22.5 μm. Because the ISIIS camera was attached to the back of the DEEP TOW, only photos taken during the down-cast were considered because the forward motion of the DEEP TOW during the up-cast could interfere with plankton moving naturally through the imaging area of the camera. A total of 4,010 photos were taken during the down-cast at station 3; 3,639 at station 10; 3,056 at station 15; and 2,453 at station 17, so that 13,158 photos were included in the study—an equivalent of 4,931.5 Liters of seawater. Down-cast photos were manually viewed by a single researcher and Regions of Interest (ROIs) containing characteristically star-shaped acantharian vegetative cells were cropped and saved. The ISIIS internal clock was calibrated to match that of a Sea-Bird SBE 9 CTD mounted to the DEEP TOW so that CTD data could be used to determine the depth at which each image was taken. Concentrations of acantharian cells per Liter were determined by normalizing acantharian cell counts to the total number of images taken for 10 m bins and correlation between cell concentration and depth was evaluated. ROI image area was used as a proxy for cell size, allowing for comparisons in cell-size range between sites and depths. Figure 2 illustrates the morphological diversity and size range of acantharians imaged in this study. Acantharian ROIs and all raw images used in the study are archived with Zenodo (https://doi.org/10.5281/zenodo.3605400).

**Fig 2.**
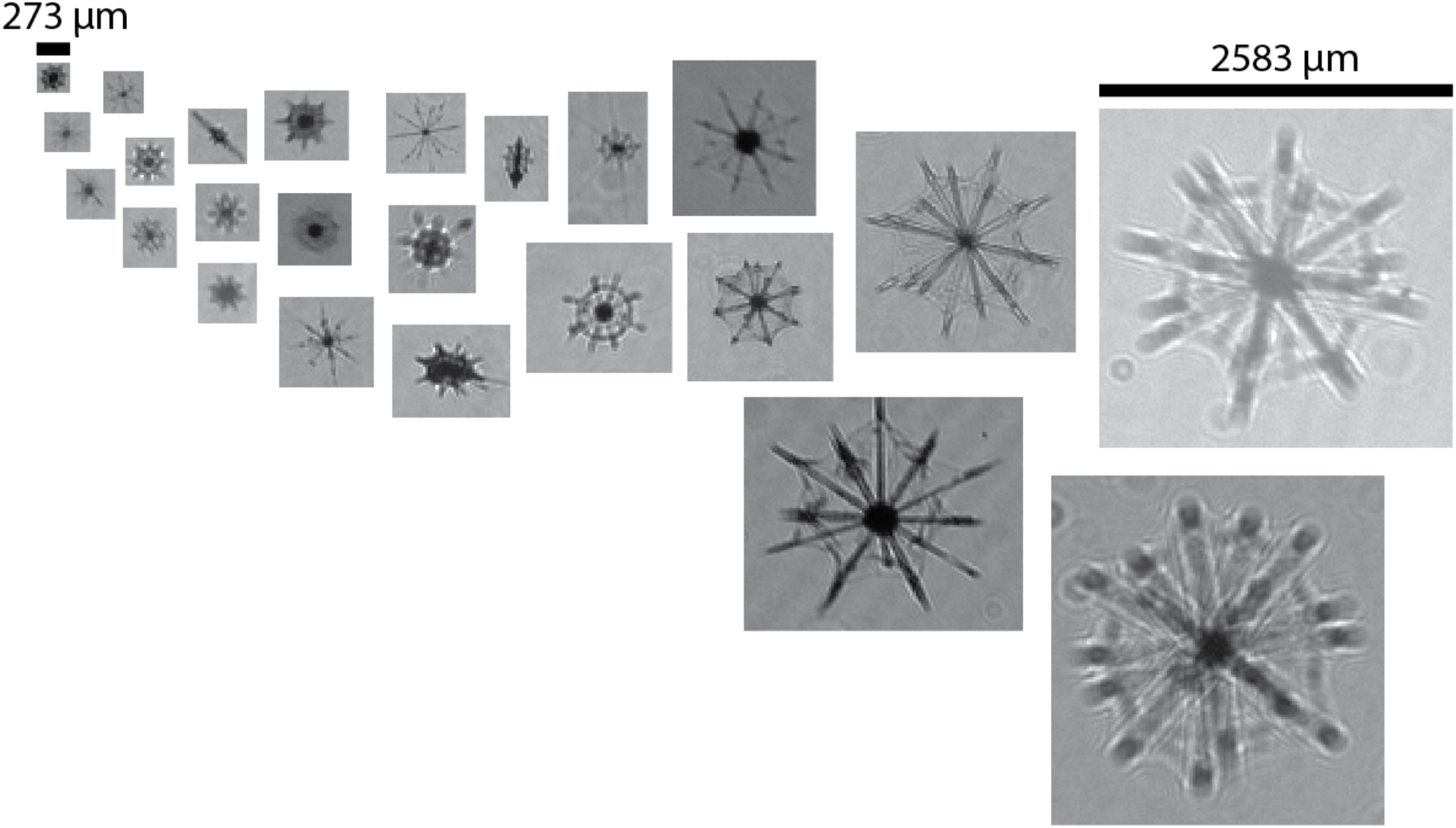
Acantharians imaged in this study, illustrating the morphological diversity and size range of imaged cells.

### Water Sampling and DNA extraction

A Niskin bottle rosette with 30 bottles (10 L) and fitted with a CTD probe (SBE 911plus, Sea-Bird Scientific, Bellevue, WA) was deployed at each cruise station to collect water from the subsurface chlorophyll maximum (SCM), the middle water column (mid), and approximately 10 m above the seafloor (bottom) (Table S1). Surface seawater was collected by bucket alongside the research vessel. Two replicates of 4.5 Liters (surface) from separate bucket casts or 5 Liters from separate Niskin bottles (SCM, mid, bottom) were sequentially filtered under a gentle vacuum through 10.0-μm and 0.2-μm pore-size polytetrafluoroethylene (PTFE) filters (Millipore, Burlington, MA). Sequential size filtering was implemented in order to separate vegetative acantharian cells and cysts from reproductive swarmer cells (< 5 μm, (Decelle et al., 2012a), although complete separation is probably not possible. Filters were flash-frozen in liquid nitrogen and stored at −80°C.

DNA was extracted from PTFE filters (n = 224, two replicates of two filter pore-sizes at four depths from 14 stations) following manufacturer’s protocols for the DNeasy PowerWater Kit (Qiagen, Hilden, Germany) including the optional heating step for 10 min at 65°C to fully lyse cells. Sequencing libraries were prepared following the Illumina 16S Metagenomic Sequencing Library Preparation manual, but with universal eukaryotic primers for the V4 region of the eukaryotic 18S rRNA gene (F: CCAGCASCYGCGGTAATTCC (Stoeck et al., 2010), R: ACTTTCGTTCTTGATYR (Mars Brisbin et al., 2018)) and 58°C annealing temperature in the initial PCR. Amplicon libraries were sequenced by the Okinawa Institute of Science and Technology (OIST) DNA Sequencing Section on the Illumina MiSeq platform with 2 × 300-bp v3 chemistry. Amplification and sequencing were successful for 211 samples, with at least one replicate for each sample type.

### Sequence analysis

Sequence data from each of four flow-cells were denoised separately using the Divisive Amplicon Denoising Algorithm (Callahan et al., 2016) through the DADA2 plug-in for QIIME 2 (Bolyen et al., 2019). Denoised Amplicon Sequence Variant (ASV) tables were merged before taxonomy was assigned to ASVs with a naive Bayes classifier trained on the Protist Ribosomal Reference (PR2) database (Guillou et al., 2013) using the QIIME 2 feature-classifier plug-in (Bokulich et al., 2018). Results were imported into the R statistical environment (R Core Team, 2018) for further processing with the Bioconductor package phyloseq (McMurdie and Holmes, 2013). Full protist communities (including all eukaryotic ASVs, except those classified as Metazoa) were analyzed first to evaluate to what degree overall community composition varied by sampling depth and by filter pore-size at each depth. Sequences classified as Acantharea were further analyzed separately to determine (i) if patterns by depth and filter pore-size for acantharians reflected overall community patterns, (ii) how much acantharian sequences contributed to the total number of sequencing reads from each depth, (iii) how the relative abundance of different acantharian clades varied by depth, and (iv) if acantharian contributions to total sequencing read numbers correlated to cell concentrations determined from imaging data. The data and code necessary to reproduce all statistical analyses are available on GitHub (https://github.com/maggimars/Acanth_ImageSeq), including an interactive online document: https://maggimars.github.io/Acanth_ImageSeq/Acanth_ImageSeq_Analysis.html.

## Results

### Sequencing Results

Overall, 31.5 million sequencing reads were generated for this study, with 34,631–421,992 sequencing reads per sample (mean = 144,604). All raw sequence data is available from the NCBI Sequencing Read Archive, accession number PRJNA546472. Following denoising, 16.8 million sequences remained and 1.1 million were classified as Acantharea. We identified 1,053 unique acantharian ASVs in our dataset, out of a total of 22,656 unique ASVs.

In Principal Coordinates Analyses (PCoA) of Aitchison distances between samples based on full community compositions, samples clustered by depth first, with clear separation of surface and SCM samples from mid and bottom water samples on the primary axis; SCM and surface samples further separated from each other on the secondary axis (Fig. S2A). When full communities were analyzed for each depth separately, surface and SCM samples separated by filter-pore size on the primary axis and mid and bottom water samples separated by filter-pore size on the secondary axis (Fig. S3A), but these results were not found statistically significant with Permutational Analyses of Variance (PERMANOVA). Notwithstanding, the clear sample clustering by filter pore-size for each depth suggests that size-fractionation was moderately successful. It remains likely, however, that larger cells were broken or otherwise squeezed through the larger pore-size filter to be captured on the lower filter, and that some smaller cells were stuck and retained on the larger pore-size filter.

When only ASVs classified as Acantharea were included in PCoA based on Aitchison distances, samples also clustered first by depth, but the overall pattern was distinct from that seen when full communities were analyzed. Acantharian communities varied more in mid and bottom water samples than full communities did (Fig. S2B). Furthermore, while the full communities clustered separately by filter pore-size in each depth layer (Fig. S3A), this was not true for acantharian communities, which did not separate by filter pore-size at any depth (Fig. S3B).

At the surface, Arthracanthida and Symphyacanthida acantharian clades made up almost the entire acantharian community in both large and small size-fraction samples at every station (Fig. 3). Arthracanthida and Symphyacanthida are the most recently diverging acantharian clades—acantharians belonging to these clades are photosymbiotic and have robust skeletons that are sometimes ornamented with elaborate apophyses (Decelle et al., 2012b). In the SCM, the contribution of sequences from Chaunacanthida acantharians increased, as did sequences deriving from the Acantharea-Group-II (Fig. 3). The Chaunacanthinda clade diverged earlier than both the Arthracanthida and Symphyacanthida clades and acantharians belonging to this clade are generally asymbiotic, have less developed skeletons, and are capable of encystment (Decelle et al., 2012b, 2013). The Acantharea-Group-II is one of several basal clades that are defined entirely by sequences recovered from environmental samples and have no known morphology (Decelle et al., 2012b). In the mid and near-bottom water, the majority of the acantharian reads derived from another basal environmental clade, the Acantharea-Group-I (Fig. 3).

**Fig. 3.**
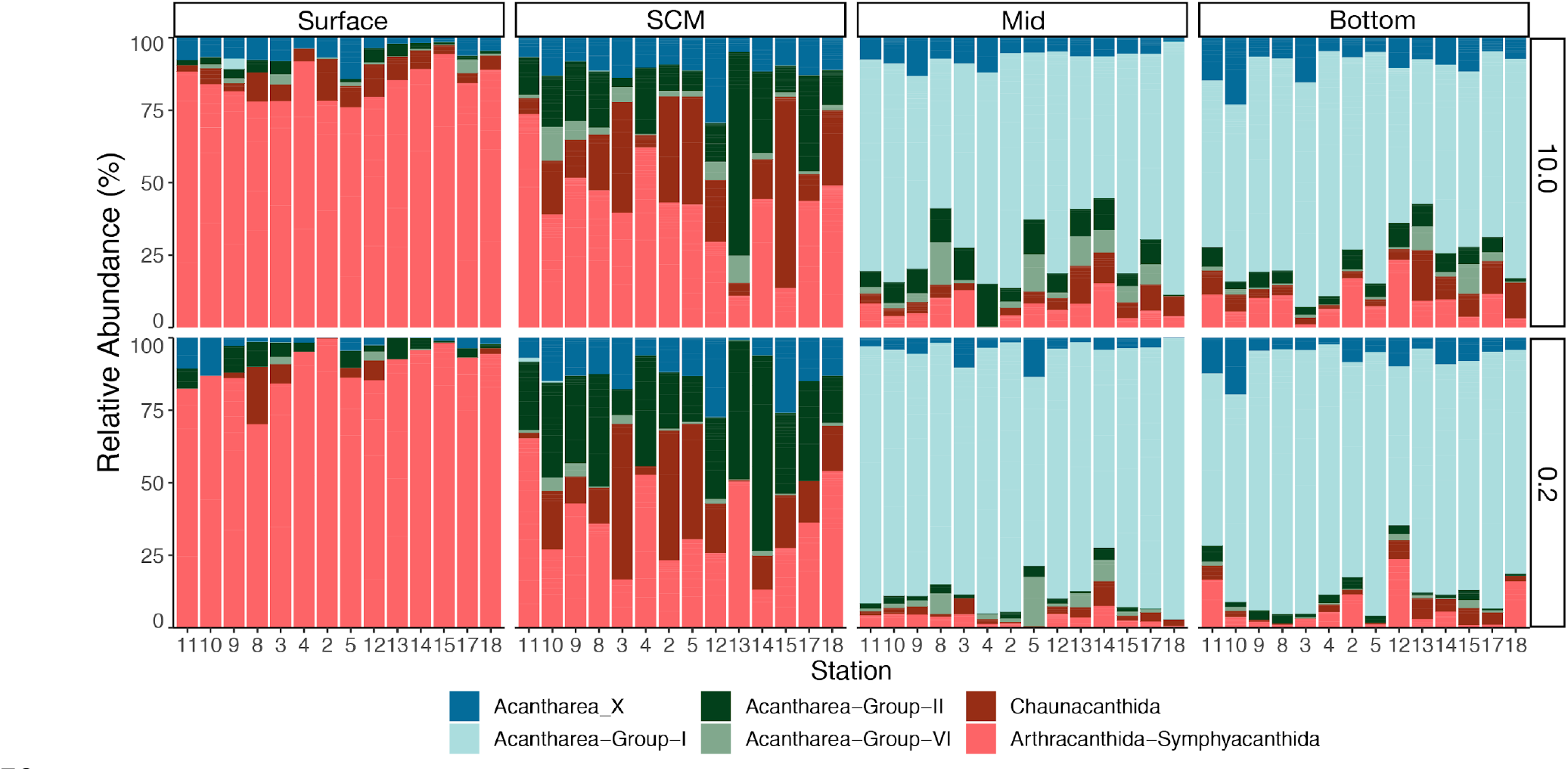
Relative abundance of acantharian groups in size-fractionated samples from four depths in the western North Pacific. Sampling stations are ordered on the x-axis from south to north and the plot is faceted by sampling depth (columns) and filter pore-size in μm (rows). When replicates were available for a particle station/depth/filter combination, replicates were collapsed and represented with a single stacked bar. Stacked bars are divided to represent the contribution of major acantharian clades to acantharian communities: dark blue (Acantharea_X) represents sequences that were not classified past the class level (Acantharea). The acantharian communities at the surface are dominated by Symphyacanthida and Arthracanthida acantharians, while at the SCM, Symphyacanthida and Arthracanthida acantharians still make up a large proportion of reads but communities are more diverse. Acantharian communities are dominated by the basal environmental clade Acantharea-Group-I in the mid and near-bottom waters.

In order to evaluate the relationship between depth and acantharian read abundance, the percent contribution of acantharian ASVs to all sequencing reads was calculated for each sample and a linear regression model was fit to the percentages with depth as the independent variable. The linear model fit to acantharian sequence percentages in all samples demonstrated a significant positive correlation to depth (R^2^ = 0.245, *p* < 0.001, Fig. 4A). To facilitate comparisons between sequencing and imaging data, linear models were also fit individually to acantharian read percentages from stations where imaging profiles were taken. Increasing acantharian sequence percentage with depth was apparent for three of the four stations with imaging profiles, but model results were only significant for station 17 (R^2^ = 0.725, *p* < 0.01, Fig. 4B).

**Fig. 4.**
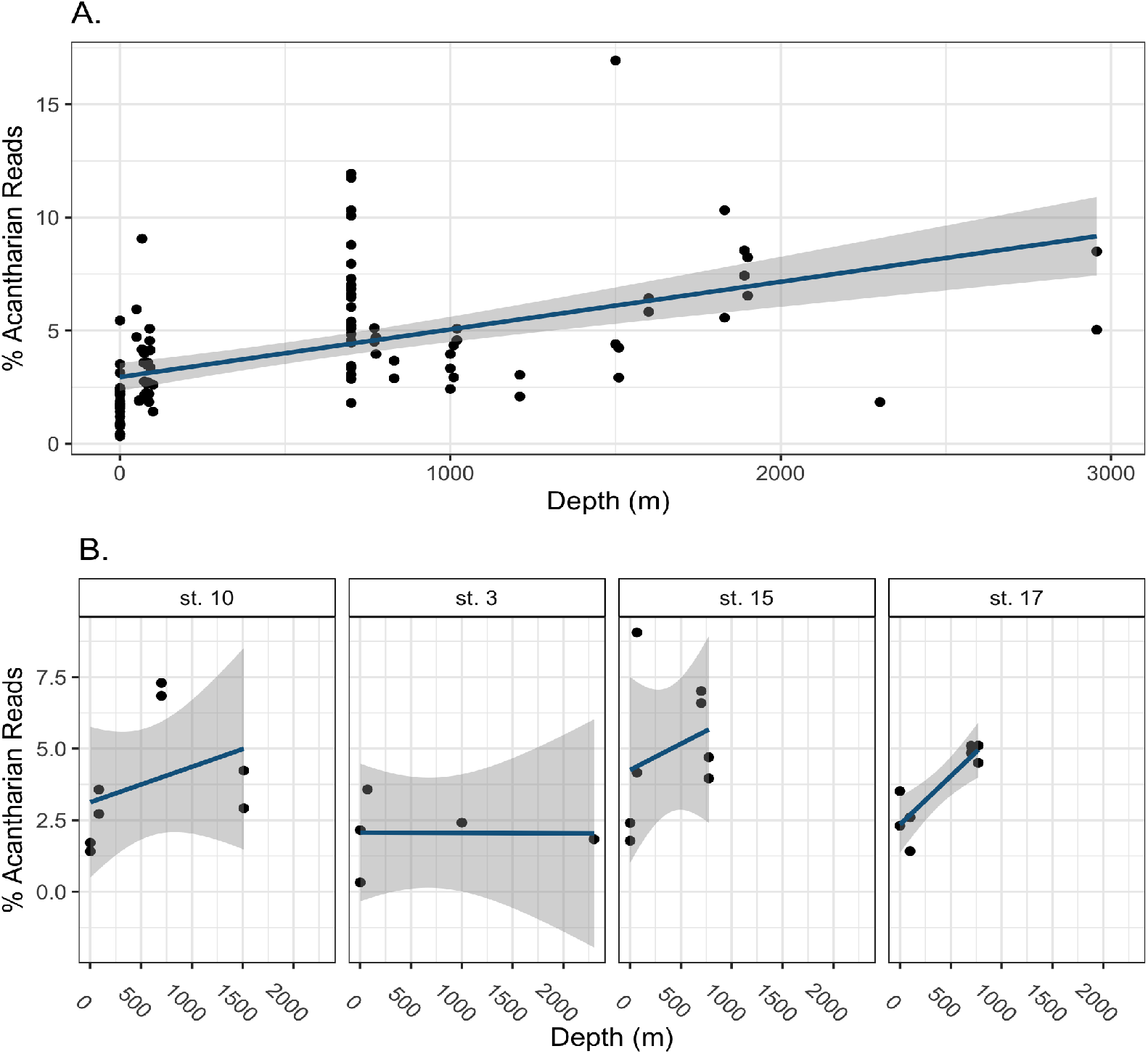
Percentage of sequences deriving from acantharians in water samples from all stations (A) and the four stations where plankton imaging was performed (B). Sequence percentages refer to the proportion of reads designated as acantharian out of all denoised sequencing reads for a given sample, including metazoan reads and reads without a taxonomic classification. Linear models were fit to the data and are represented by blue lines with the 95% confidence intervals shaded grey. The linear model results for all samples (A) are statistically significant (*p* < 0.001) with an R^2^ of 0.2516. The linear model results for individual stations (B) were only statistically significant for station 17 (*p* < 0.01) with an R^2^ of 0.76. Overall, the contribution of acantharians to the whole community was smallest in the surface waters and tended to increase with depth sampled.

Only acantharians in the Chaunacanthida, Arthracanthida, and Symphyacanthida clades are known to possess the characteristic star-shape used to identify acantharian vegetative cells in images from this study. Therefore, the percentages of Chaunacanthida, Arthracanthida, and Symphyacanthida were further considered separately. When only percentages for these clades with known morphologies were included in the model, read abundance was significantly negatively correlated with depth (R^2^ = 0.170, *p* < 0.001, Fig. 5A)—opposite to the relationship when all clades were analyzed together. The same trend was apparent for all stations with imaging profiles when evaluated individually—percentage decreased with depth—but model results were only statistically significant for station 10, with R^2^ = 0.65 and *p* < 0.01 (Fig. 5B).

**Fig. 5.**
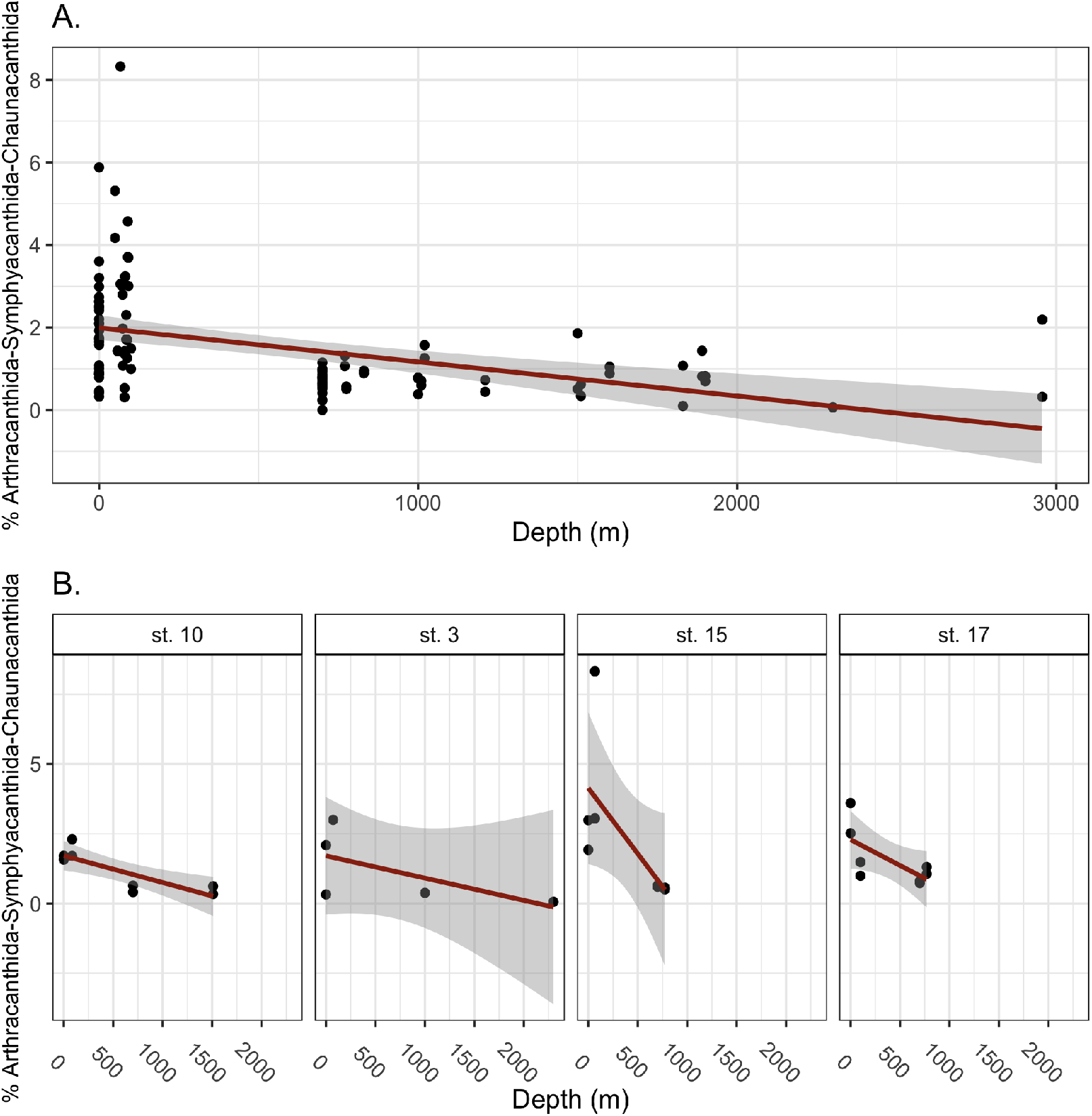
Percentage of sequencing reads deriving from Symphyacanthida, Arthracanthida, and Chaunacanthida acantharians in water samples from all stations (A) and the four stations where plankton imaging was performed (B). Percentages refer to the proportion of reads designated as Symphyacanthida, Arthracanthida, or Chaunacanthida acantharians out of all denoised sequences for a given sample, including metazoan sequences and sequences without a taxonomic classification. Linear models were fit to the data and are represented by red lines with 95% confidence intervals shaded grey. The linear model results for all samples (A) were statistically significant (*p* < 0.001) with an R^2^ of 0.1697. The linear model results for individual stations (B) were not statistically significant (R^2^ = 0.273–0.653). Overall, the contribution of Symphyacanthida, Arthracanthida, and Chaunacanthida acantharians to the whole community was larger in surface waters and at the SCM and tended to be smaller in the mid and near-bottom waters.

### Imaging Results

Overall, 1,235 acantharian ROIs were identified in the study and the vast majority of these were found in images taken near the sea surface (Fig. 6A). When linear models were fit to log-transformed cell concentrations calculated from images, with depth as the independent variable, cell concentrations were significantly negatively correlated with depth at each station (R^2^ = 0.631–0.777, *p* < 0.001, Fig. 6A). These results match the relationship between Chaunacanthida, Arthracanthida and Symphyacanthida sequence percentages with depth but not the relationship between all acantharian sequence percentages with depth. To directly evaluate how well imaging results correlated with sequencing results, we averaged acantharian cell concentrations in each depth layer (surface: 0–50 m, SCM: 50–150 m, mid: 150–700 m, deep/bottom: > 700 m) and compared these values to Chaunacanthida, Arthracanthida and Symphyacanthida sequence percentages in samples from corresponding stations and depths. Averaged cell concentrations significantly positively correlated with Chaunacanthida, Arthracanthida and Symphyacanthida relative sequencing read abundance (R^2^ = 0.334, *p* < 0.05) following exclusion of two outlying data points with exceptionally high sequencing read abundance or imaged cell concentration (Fig. 7). Acantharian cell size ranged widely in the surface waters and near the SCM, whereas the range was more constrained and cell size was generally smaller in deeper water (Fig. 6B). Interestingly, many acantharians were observed with long pseudopodial extensions terminating in drop-shaped structures (Fig, 8). This morphology has not previously been observed in acantharians, probably due to damage caused by plankton nets or other handling effects.

**Fig. 6.**
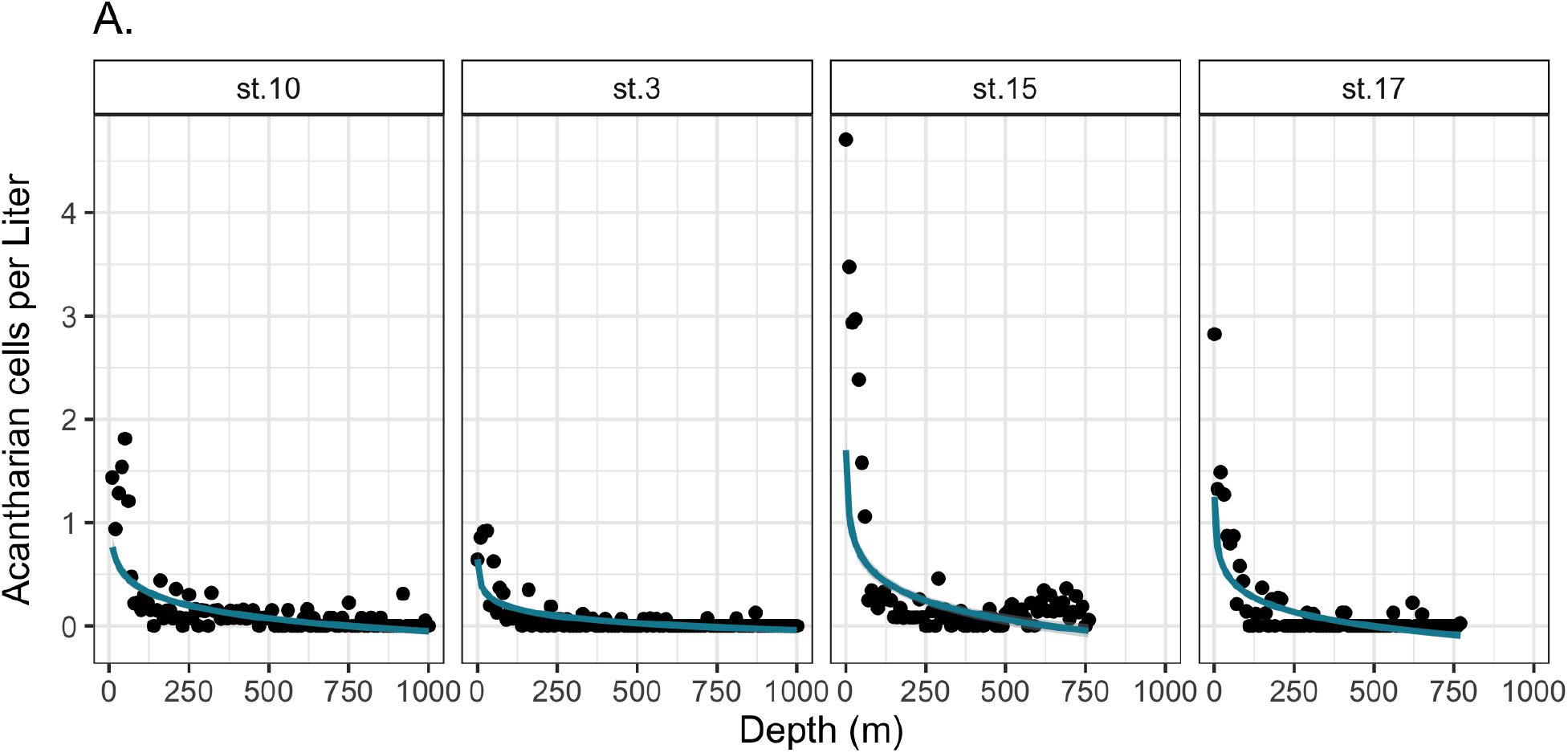
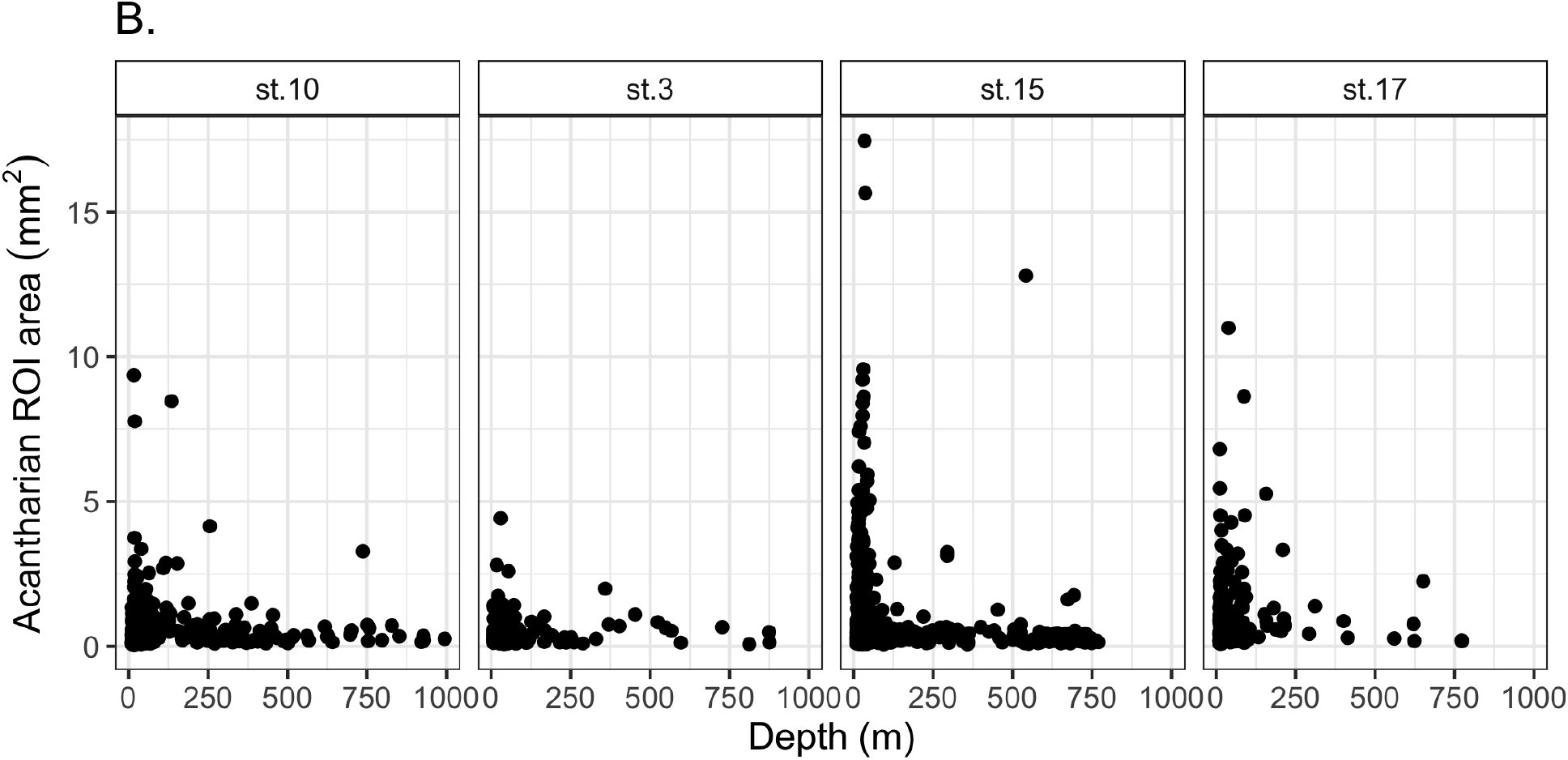
Concentrations of visible acantharian cells per Liter observed in vertical imaging profiles and size distributions of acantharian Regions of Interest (ROIs) (B). Acantharian cell concentrations were determined by dividing the number of observed cells for a 10 m section of the vertical profile by the volume of water imaged in that section. Linear models were fit to log-transformed data and are plotted in turquoise with 95% confidence intervals shaded grey (visible for station 15). Linear models for each station (A) were statistically significant (*p* < 0.001) with st. 10 R^2^ = 0.64; st. 3 R^2^ = 0.63; st. 15 R^2^ = 0.66; st. 17. R^2^ = 0.78. The highest concentrations of visible acantharian cells were always observed close to the sea surface and decreased sharply with depth. Acantharian ROIs were cropped so that the edges of the rectangular photos aligned with the outward reaches of cellular extensions in each direction. The pixel dimensions of each ROI image were converted to microns and then the area of ROI images was calculated and plotted against the depth at which it was imaged (B). ROIs exhibit a large size range in surface waters but larger ROIs become less common in deeper waters.

**Fig. 7.**
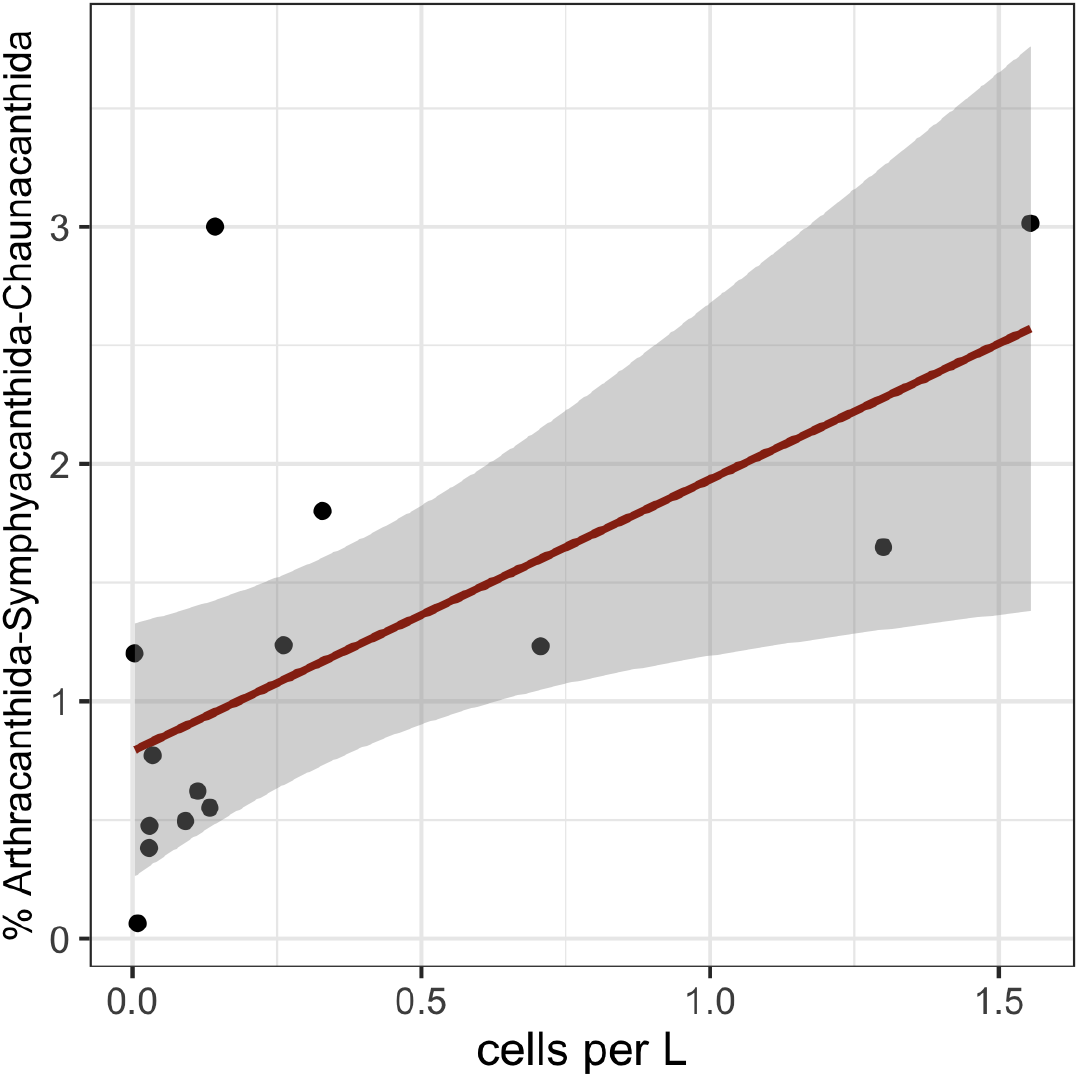
Linear regression of acantharian cells per Liter averaged for surface, SCM, mid, and bottom depth layers against percent read contributions of Arthracanthida, Symphyacanthida, and Chaunacanthida acantharians at stations 2, 10, 15 and 17. Acantharian cell concentration determined by high-throughput, in-situ imaging significantly correlates with sequencing read percentages of acantharians with known morphologies (R^2^ = 0.334, *p* < 0.05).

**Fig. 8.**
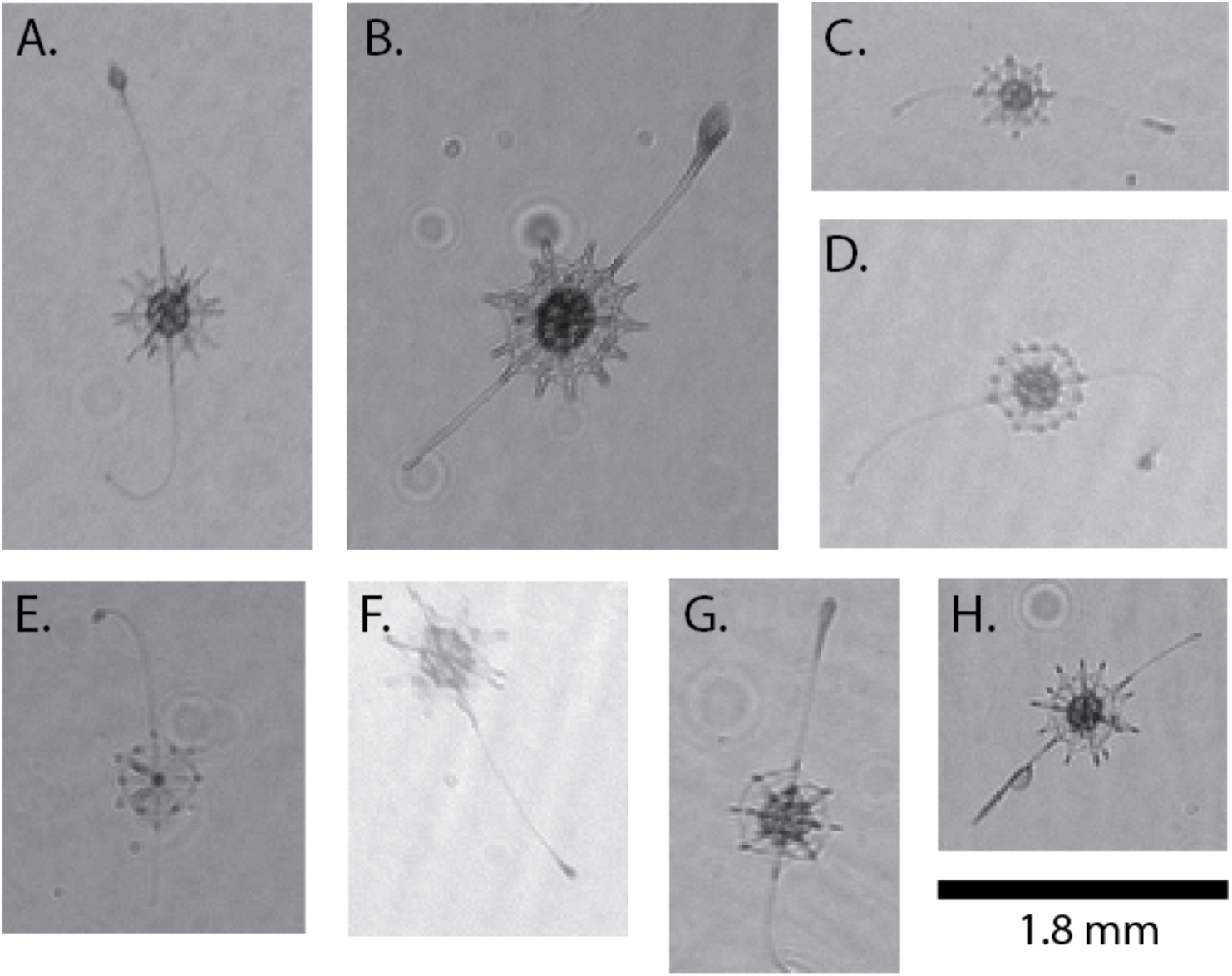
In-situ imaging reveals potential acantharian predation behavior. Acantharians imaged in this study were observed with long pseudopodial extensions ending in a drop-shaped structures. This morphology has not been previously observed, probably because the extensions are damaged in net-collected samples. Images were taken at 13.8 m, st. 15 (A); 16.4 m, st. 15 (B, C, E); 27.5 m, st. 15 (D);31.2 m, st. 15 (F); 57.2 m, st. 17 (G); 13.8 m, st. 17 (H). Image aspect ratios are unaltered and the scale bar below panel H is accurate for all panels. Image orientations are likewise unchanged, with the top of images being toward the sea surface.

## Discussion

Acantharians are important contributors to primary production throughout the global ocean, but detailed studies on absolute abundance and fine-scale distribution have been hindered by specific acantharian traits, such as their fragile cell structures and skeletons that dissolve in common fixatives. In addition, the smaller size of acantharians compared to other Rhizarians has precluded their full inclusion in quantitative imaging surveys (Biard et al., 2016; Biard and Ohman, 2019). As a result, advances in our understanding of acantharian biology and ecology have come primarily from molecular studies. In this study, we took a paired approach and combined molecular survey methods with high-throughput, in-situ imaging to better evaluate acantharian abundance and distribution. We found that vegetative acantharian cells were concentrated in the uppermost water column, but were sporadically present throughout the water column, including at the deepest depths images were taken (1,000 m, Fig. 6B). The concentrations of acantharian cells determined from imaging data correlated well with the contribution of Arthracanthida, Symphyacanthida, and Chaunacanthida acantharians to sequences recovered from the same depth and location (Fig. 7). In contrast, the percentages of sequences from all acantharians, including those belonging to undescribed environmental clades, increased with depth. Taken together, the results can provide information about the distribution and abundance of different clades of acantharians, the morphology of undescribed environmental clades, and the life cycles of acantharians.

### Acantharian abundance and distribution

In this study, we observed maximum acantharian abundances of 0.9–4.7 cells per L using an in-situ camera system capable of imaging organisms with diameters greater than 250 μm. Maximum acantharian abundances were observed in the upper euphotic zone at each station: 0–10 m depth at stations 15 and 17, 20–30 m depth at station 3, and 40–50 m depth at station 10 (Fig. 6A). These results are consistent with previous studies that carefully preserved and counted acantharians collected by high-volume plankton pump and with Niskin bottles (Michaels, 1991; Michaels et al., 1995). Michaels et al. (1995) observed near-surface acantharia maxima in the subtropical North Atlantic with maximum abundances ranging from 5.5–18 cells per L (mean 1.2 cells with > 100 μm diameter per L in Niskin samples; mean 2.5 cells per L in pumped samples). Similarly, Michaels (1991) recorded 0.1–4 acantharian cells per L (> 100 μm diameter) in the surface mixed layer of the eastern North Pacific Subtropical Gyre. Using high-throughput, in-situ imaging, Biard and Ohman (2016) likewise found high acantharian concentrations near the surface in the California Current Ecosystem, although they identified acantharians only with diameters > 600 μm. Compared to other high-throughput imaging studies (Biard et al., 2016; Biard and Ohman, 2019), the acantharian abundances we recorded are closest to the cell abundances reported when cells were counted by microscopy. While the cell abundances we measured are likely still an underestimate, since many acantharians are smaller than 250 μm (Michaels, 1991; Michaels et al., 1995), the results from our high-throughput imaging allow for a more quantitative estimate of acantharian abundance and vertical distribution in the western North Pacific than analyzing sequencing data alone.

### Basal environmental clades of Acantharea

The relative abundance of sequences classified as Chaunacanthida, Symphyacanthida and Arthracanthida decreased as sampling depth increased (Fig. 5), which correlated with the acantharian abundances determined from imaging data (Fig. 7). In contrast, the relative abundance of sequences classified as Acantharea at the class level increased with depth and did not correlate with imaging results. The additional acantharian sequences in communities from deeper water were primarily classified as Acantharea-Group-I (Fig. 3), which is basal to Chaunacanthida, Arthracanthida and Symphyacanthida, and has no known morphology (Decelle et al., 2012b; Decelle and Not, 2015). Similarly, the acantharian contribution to clone libraries from coastal waters near California increased with depth (Schnetzer et al., 2011) and environmental sequences from basal acantharian clades I–III have been recovered from deep waters throughout the global ocean (López-García et al., 2001; Edgcomb et al., 2002; Countway et al., 2007; Not et al., 2007; Terrado et al., 2009; Gilg et al., 2010; Quaiser et al., 2011; Decelle et al., 2013). Here, the discrepancy between the depth-related increase in read abundance for Acantharea-Group-I and the coinciding decline in cells imaged with characteristic acantharian morphologies may provide new evidence regarding the morphology of basal environmental acantharian clades.

The acantharian skeleton is a central feature to their morphological classification; the most recently diverged clades (Arthacanthida and Symphyacanthida) have spicules of varying lengths—some with elaborate protrusions—that are fused in robust central junctions, whereas earlier diverging clades (e.g. Chaunacanthida) have simpler spicules of equal length that either cross the central region of the cell or form loosely-fused central junctions (Decelle et al., 2012b). This evolutionary trajectory—from less to more developed skeletons—suggests that the earliest diverging acantharian clades (i.e. basal environmental clades I–III) may have only rudimentary skeletal structures, or may lack the quintessential acantharian skeleton altogether (Decelle et al., 2012b). The decreased observance of recognizable acantharian cells with depth coinciding with an increased read abundance for clade I acantharians suggests they may, indeed, lack traditionally recognized acantharian morphologies. Alternatively, Group I acantharians may simply be too small to be imaged with the ISIIS small-imager used in this study. Ultimately, the morphology of the basal environmental acantharian clades can only be definitively resolved with single-cell sequencing of deep-sea isolates coupled with microscopy (Sieracki et al., 2019). However, our results demonstrate that small cells lacking symmetrical strontium sulfate skeletons should be considered for sequencing in studies seeking to determine the morphology of the earliest diverging acantharian clades.

### Acantharian life cycles

Knowledge regarding acantharian life cycles remains relatively limited because a full acantharian life cycle has not yet been observed. However, cyst formation and swarmer release from cysts and vegetative cells have been observed in laboratory settings (Decelle et al., 2012a, 2013). Swarmers are small (2–3 μm) biflagellated cells with unknown ploidy; it is assumed that they are reproductive cells and fuse to form juveniles, but this has never been witnessed (Decelle et al., 2012a). So far, cyst formation has only been observed for earlier diverging acantharian lineages and has not been observed for Arthracanthida or Symphyacanthida acantharians. Acantharians that form cysts shed most of their spicules before cyst formation, suggesting that acantharians in later diverging clades with more robust and elaborate skeletons cannot form cysts because of the fixed central junctions in their skeletons. In addition, cysts recovered from sediment traps have only ever been of earlier diverging clades based on phylogenetic analysis—but not environmental clades I or II (Decelle et al., 2013). As a result, current hypotheses propose that acantharians in earlier diverging clades, including Chaunacanthida, form cysts as a means for ballast, allowing them to sink to deep water where they release swarmer cells, whereas acantharians in the later diverging Arthracanthida and Symphyacanthida clades complete their life cycle in the photic zone, since they cannot form cysts and need to acquire photosymbionts at the start of each generation (Martin et al., 2010; Decelle et al., 2013).

The imaging results demonstrating decreased cell size and abundance below the surface mixed layer (Fig. 6B) are consistent with the idea that many acantharians sink to release swarmers with juveniles growing in size as they make their way up towards the surface. Adult cells sink quickly, whether for reproduction or as detritus (Gutierrez-Rodriguez et al., 2019), and both vegetative cells and cysts dissociate after releasing swarmers (Decelle et al., 2012a, 2013), making them less likely to be caught on camera. A large number of swarmers released at depth would potentially produce many small juvenile cells that gradually increase in size as they slowly ascend. These smaller, more abundant juveniles would be much more likely to be imaged than the rarer, faster moving adults. Alternatively, decreased cell-size range at depth could also reflect decreased nutritional resources available in deeper waters or constitutively smaller sized species being more common below the surface mixed layer. However, the sporadic presence of large cells in very deep water is evidence that they at least occasionally reach deep water from the surface or that cells can reach larger sizes at depth (Fig. 6B).

By combining metabarcoding of size-fractionated samples with imaging, further insight into acantharian life cycles can be gained. In principle, vegetative cells and cysts should have been retained on the upper filter with larger pore-size and the swarmer cells should have passed through the upper filter and been retained on the lower filter with smaller pore-size. A disparity in the contribution of one clade to sequences in the two size-fractions could, therefore, indicate that vegetative cells or reproductive cells from that clade are more or less abundant at a particular depth. Such size separation can never be perfect—and it may be especially problematic with delicate cells like acantharians—but Principal Coordinate Analysis (PCoA) of Aitchison distances between full protistan communities showed clear segregation by filter type at each depth (Fig. S3A), suggesting size-fractionation was relatively successful. In contrast, PCoA for acantharian communities in the same samples did not show segregation by filter type (Fig. S3B), thus providing some evidence that swarmers and vegetative cells or cysts coexist in the four depth layers sampled. Given that acantharians belonging to the Chaunacanthida clade are among those that form cysts and are believed to sink before releasing swarmers, Chaunacanthida would be expected to be more abundant than non-cysting Arthracanthida and Symphyacanthida in both size fractions at depth. Interestingly, Arthracanthida and Symphyacanthida sequences were recovered from both large and small size-fraction samples from mid and near-bottom water at every sampling station and had similar relative abundances to Chaunacanthida sequences (Fig.3). This lack of differentiation in deep water sequence abundances of Arthracanthida-Symphyacanthida and Chaunacanthida does not support the idea that the later diverging clades remain in the photic zone to reproduce. An alternative hypothesis might be that Arthracanthida and Symphyacanthida also sink to deep water to reproduce but do so as vegetative cells, aided by their robust skeletons and fine buoyancy control (Febvre and Febvre-Chevalier, 2001), rather than in the form of cysts (Michaels et al., 1995). It cannot be ruled out that the DNA recovered from deep waters could be extracellular or derive from detrital matter, but the alternative hypothesis is further supported by the occasional observation of large acantharian cells at depth (Fig. 6B, (Biard and Ohman, 2019).

### Acantharian behavior revealed by in-situ imaging

Being notoriously delicate and sticky, acantharians are often broken or clumped when collected by plankton net. As a result, their fine structure is usually damaged even when they do survive collection, which can preclude behavioral observations. In-situ imaging is especially useful in such cases, as it allows for the observation of natural orientation and behaviors that could not otherwise be seen. Acantharians are known to be active predators: microscopy of SCUBA collected acantharians revealed ciliates, diatoms, and dinoflagellates as acantharian prey items (Swanberg and Caron, 1991) and results from 18S sequence analysis of single acantharians included copepod, diatom, and dinoflagellate sequences (Mars Brisbin et al., 2018). However, actual predatory strategies of acantharians are unknown. In this study, we repeatedly observed acantharians that had long pseudopodial extensions terminating in a drop-shaped structures. This morphology/behavior has not been previously observed and we hypothesize that the extensions may represent a fishing apparatus that allow acantharians to lure and capture prey. However, since we did not observe prey items stuck to the droplets, it remains possible that these structures may be involved in other processes (e.g. reproduction, buoyancy, or locomotion).

### Prospects for future automatic classification of acantharians

The images produced in this study were annotated manually, which represents a major barrier in high-throughput imaging studies; manual image annotation is a large time commitment and requires experience identifying plankton groups. The ultimate goal for high-throughput imaging is to have annotated training sets that are extensive and comprehensive enough to provide highly accurate automatic image classification using machine learning algorithms. Currently, several instrument- and location-specific learning sets are available in the public realm: e.g. the WHOI-plankton dataset, which includes 3.4 million annotated images in 70 classes that were taken with the Imaging Flow Cytobot in Martha’s Vineyard (Orenstein et al., 2015), and the PlanktonSet-1.0, which includes 30,336 images in 121 classes that were taken with the ISIIS line-scan imager in the Straits of Florida (Rodrigues et al., 2018). These datasets could be used for transfer learning to improve classification accuracy and efficiency, but since they do not include very many acantharian images and the plankton size range for both data sets excludes the majority of acantharians imaged in this study (Rodrigues et al., 2018), they alone would not have allowed for accurate classification of acantharians in our dataset. Therefore, by contributing over a thousand new annotated acantharian images, including cells ranging in diameter from 250–2500 μm, this study will facilitate accurate automatic acantharian classification in future datasets acquired with the ISIIS small-imager and other imaging systems that capture a similar size-range of organisms. Accurate automatic classification will eventually allow for larger studies of acantharian abundance and distribution, including expanded geographic and temporal scales, and thus a deeper understanding of acantharian contributions to biogeochemical processes in the ocean.

## Conclusions

High-throughput imaging used in this study showed that acantharians are abundant in the surface waters of the western North Pacific (East China Sea) and have similar per Liter concentrations as have been reported in the eastern North Pacific and the North Atlantic where cells were manually counted with microscopy. Similar to previous studies, vegetative acantharian cells were concentrated very close to the sea surface and decreased in abundance with depth, but were still sometimes observed at depths approaching 1000 m. Imaging data correlated with sequence abundances from acantharian clades with known and easily recognizable morphologies, but were in contrast to sequence abundances from acantharian environmental clade I, whose morphology is not known. This discrepancy suggests that basal environmental clades, such as clade I, may have morphologies distinct from other acantharians and may lack characteristic star-shaped strontium sulfate skeletons. The size distribution of imaged acantharians is consistent with current hypotheses about acantharian life cycles: size range decreases with depth, supporting the idea of reproduction at depth with small swarmer cells, followed by the ascension, and growth, of juveniles into surface waters. However, the similar relative abundance of different acantharian clades in small and large size fractions at depth suggests that later diverging clades (i.e. Arthracanthida and Symphyacanthida) may also reproduce at depth, which is counter to previous hypotheses. By pairing high-throughput sequencing with high-throughput, in-situ imaging, this study advances our understanding of acantharian biology but also highlights how much is still unknown. Future studies will benefit from the annotated images produced in this study, but should consider further pairing imaging with RNA sequencing or single-cell genomics.

## Acknowledgments

We thank the captain and crew of the JAMSTEC R/V *Mirai* for their assistance and support in sample collection. Hiroyuki Yamamoto, Hiromi Watanabe, Dhugal Lindsay, and Yuko Hasagawa were instrumental in organizing and facilitating cruise sampling. Dhugal Lindsay, Andrew Carroll, and Mehul Sangekar deployed the DEEP TOW and managed imaging systems. We thank the OIST DNA sequencing section (Onna, Okinawa) for carrying out the sequencing. This work was funded by the Marine Biophysics Unit of the Okinawa Institute of Science and Technology Graduate University. MMB was supported by a Japan Society for the Promotion of Science DC1 graduate student fellowship.

## Supplementary Materials

**Table S1.** Coordinates, sampling depths, and total depth for all sampling stations.

| Station | Longitude (°E) | Latitude (°N) | “SCM” Depth (m) | “Mid” Depth (m) | “Bottom” Depth (m) | Site Depth (m) |
| --- | --- | --- | --- | --- | --- | --- |
| 2 | 126.468 | 26.290 | 92 | 700 | 1066 | 1080 |
| 3 | 127.500 | 25.416 | 72 | 1000 | 2385 | 2407 |
| 4 | 126.900 | 25.928 | 58 | 700 | 1834 | 1857 |
| 5 | 126.084 | 26.501 | 82 | 700 | 1900 | 1922 |
| 8 | 124.994 | 25.942 | 88 | 700 | 1671 | 1681 |
| 9 | 124.012 | 25.502 | 74 | 700 | 1890 | 1914 |
| 10 | 123.837 | 24.857 | 85 | 700 | 1515 | 1530 |
| 11 | 122.840 | 24.760 | 50 | 700 | 1217 | 1223 |
| 12 | 126.903 | 27.785 | 80 | 700 | 1024 | 1033 |
| 13 | 127.339 | 29.003 | 80 | 700 | 834 | 846 |
| 14 | 127.501 | 28.747 | 75 | 700 | 1013 | 1025 |
| 15 | 129.029 | 28.792 | 67 | 700 | 776 | 783 |
| 17 | 129.570 | 28.957 | 90 | 700 | 772 | 779 |
| 18 | 130.904 | 28.981 | 100 | 1500 | 2957 | 2981 |

**Fig. S1.**
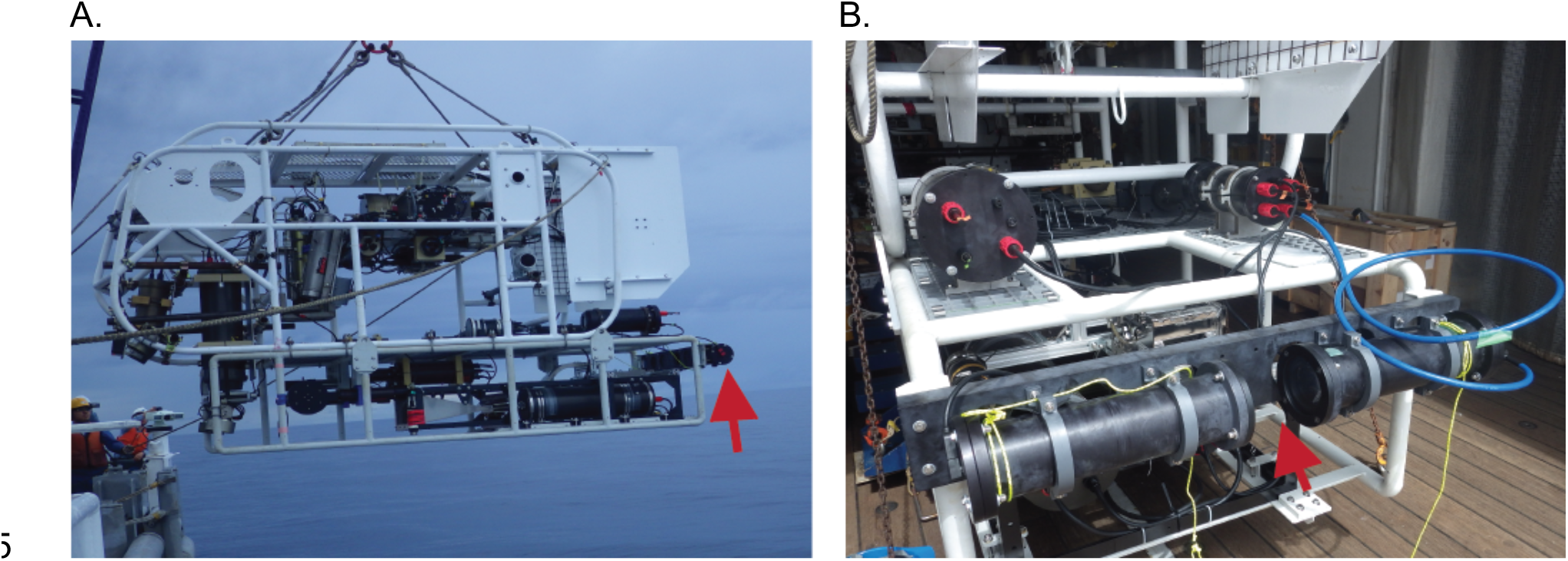
Photos of the JAMSTEC DEEP TOW 6KCTD (A) and BellaMare ISIIS small imager/area scanner attached to the DEEP TOW (B). Red arrows indicate the position of the ISIIS system on the DEEP TOW (A) and the imaging area of the ISIIS system (B). At each sampling station, the DEEP TOW was lowered straight down through the water column, to a maximum depth of 1000 m, before being towed at 1000 m and then continuing to be towed as it was raised back up through the water column. Only images from downward casts were used for this study, since forward motion of the frame could prevent plankton from being properly imaged by the ISIIS.

**Fig. S2.**
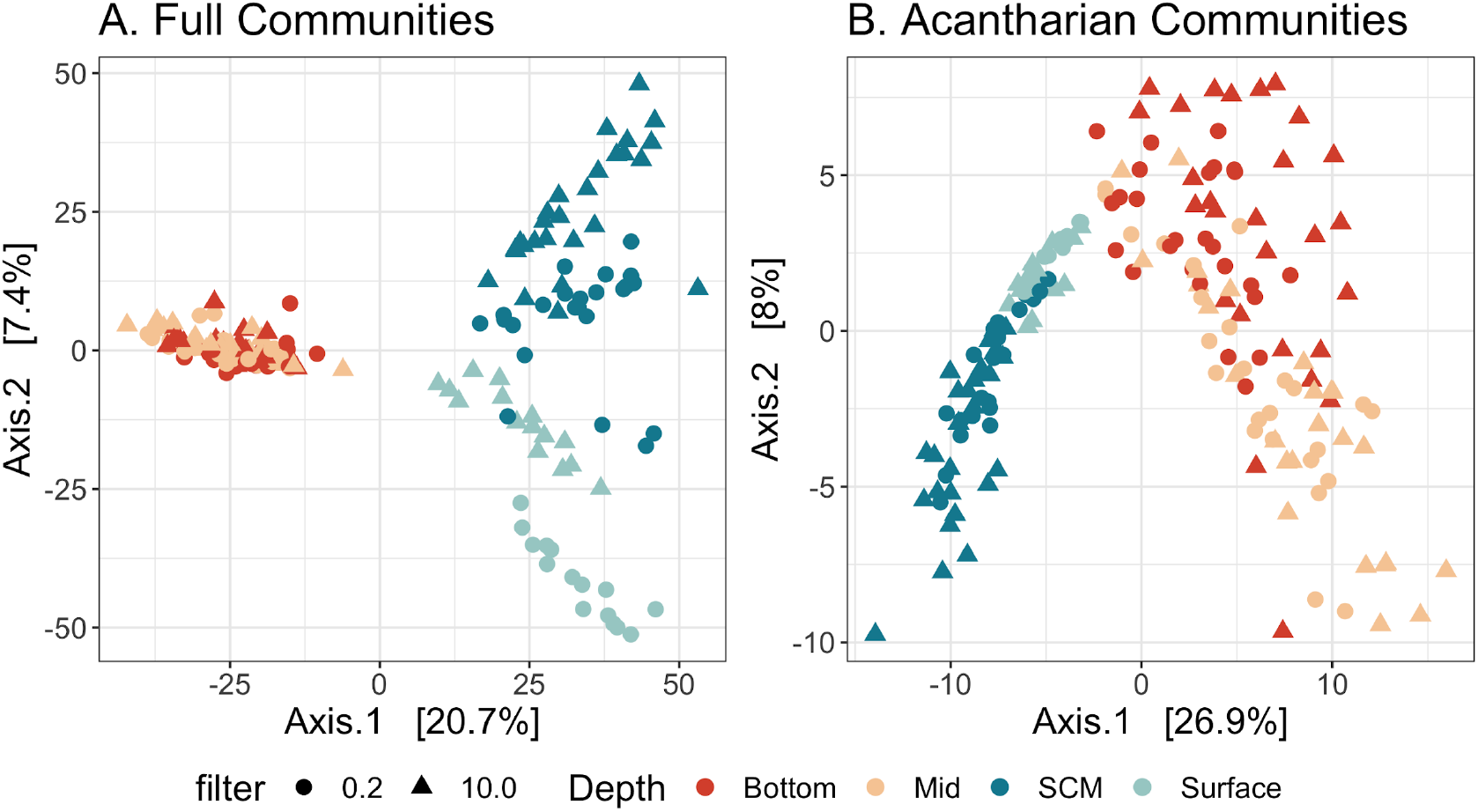
Principal coordinates analysis of Aitchison distance between full protistan community compositions (A) and acantharian community compositions (B). Full communities (A) include all denoised protist sequences that were classified as Eukaryota, but not Metazoa. Acantharian communities (B) include all denoised sequences classified as Acantharea in the 4th taxonomic level of the PR2 database. Color indicates the depth layer from which samples were collected and shape reflects the filter pore-size used to collect samples in μm. Full communities form three main clusters by depth—surface, subsurface chlorophyll maximum (SCM), and mid/bottom (A)—whereas the acantharian communities are more varied in the mid and near-bottom waters (B).

**Fig. S3.**
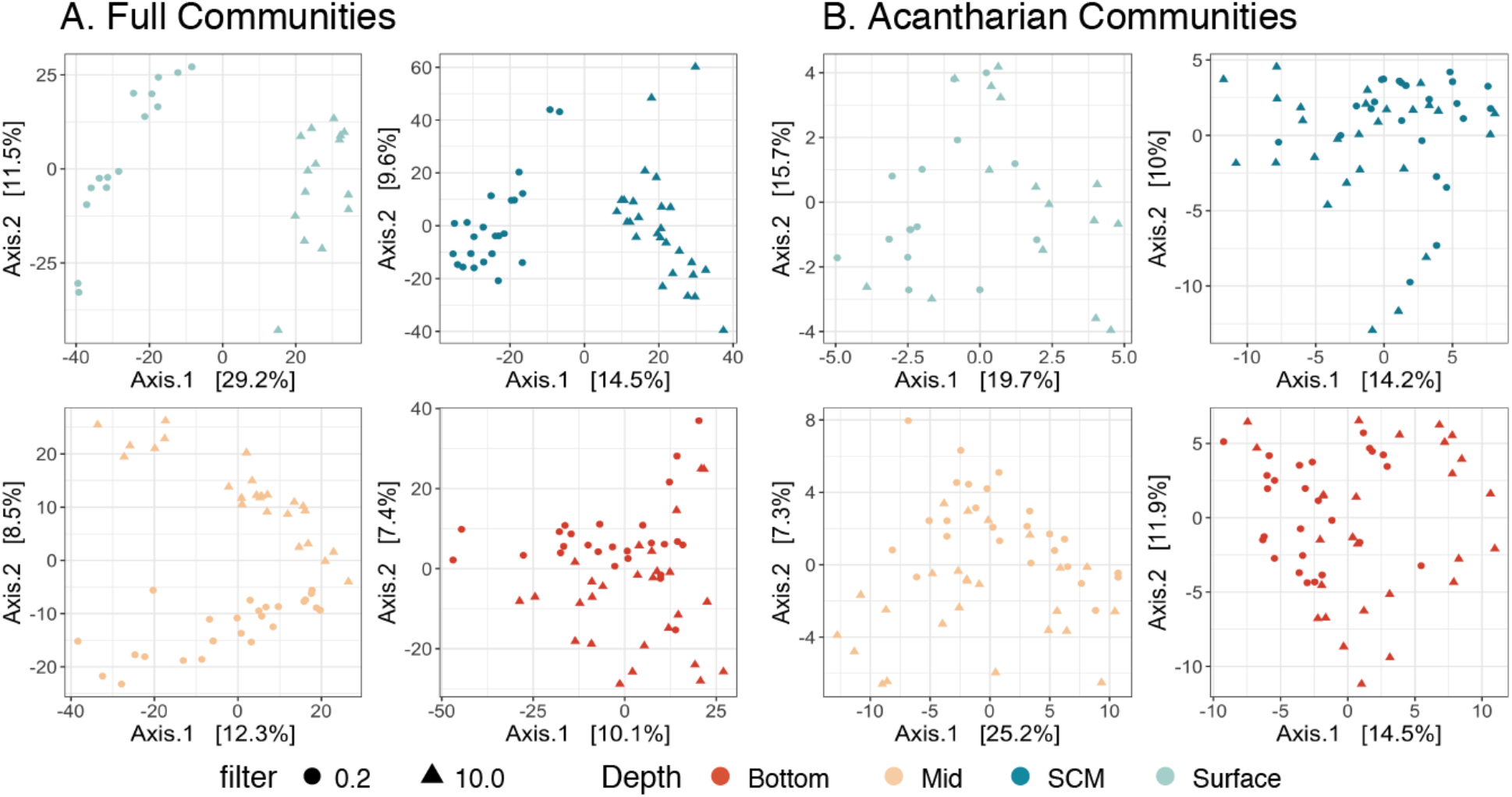
Principal coordinates analysis of Aitchison distances between eukaryotic plankton community compositions (A) and acantharian community compositions (B) in different depth layers. Separating samples by depth layer allows for better resolution of the effect of filter pore-size on community composition. Color indicates the depth layer from which samples were collected and shape reflects the filter pore-size used to collect samples in μm. Full communities (A) include all denoised protist sequences that were classified as Eukaryota, but not Metazoa. Acantharian communities (B) include all denoised sequences classified as Acantharea in the 4th taxonomic level of the PR2 database. Full communities cluster by filter pore-size along the primary axis at the surface and SCM and along the secondary axis in mid and near-bottom waters (A); acantharian communities do not cluster by filter pore-size at any sampling depth (B).

